# Binge-like Acquisition of α-pyrrolidinopentiophenone (α-PVP) Self-Administration in Female Rats

**DOI:** 10.1101/191155

**Authors:** Mehrak Javadi-Paydar, Eric L. Harvey, Yanabel Grant, Sophia A. Vandewater, Kevin M. Creehan, Jacques D. Nguyen, Tobin J. Dickerson, Michael A. Taffe

## Abstract

The synthetic cathinone α-pyrrolidinopentiophenone (α-PVP) has been associated with violent and/or bizarre public behavior in users. Association of such behavior with extended binges of drug use motivates additional investigation, particularly since a prior study found that half of male rats experience a binge of exceptionally high intake, followed by sustained lower levels of self-administration during the acquisition of intravenous self-administration (IVSA) of a closely related drug, 3,4-methylenedioxypyrovalerone. The binge-like acquisition pattern appeared to be novel for rat IVSA, thus the present study was designed to determine if this effect generalizes to IVSA of α-PVP in female rats. Female Wistar rats were trained in IVSA of α-PVP (0.05 mg/kg/inf) in experimental chambers that contained an activity wheel. Groups of animals were trained with the wheels fixed (No-Wheel group), fixed for the initial 5 days of acquisition or free to move throughout acquisition (Wheel group). The groups were next subjected to a wheel-access switch and then all animals to dose-substitution (0.0125-0.3 mg/kg/inf) with the wheels alternately fixed and free to move. Approximately half of the rats initiated their IVSA pattern with a binge day of exceptionally high levels of drug intake, independent of wheel access condition. Wheel activity was much lower in the No-Wheel group in the wheel switch post-acquisition. Dose-effect curves were similar for wheel-access training groups, for binge/no binge phenotypic subgroups and were not altered with wheel access during the dose-substitution. This confirms the high reinforcer efficacy of α-PVP in female rats and the accompanying devaluation of wheel activity as a naturalistic reward.

## Introduction

The structurally related cathinone derivative psychostimulant drugs 3,4-methylenedioxypyrovalerone (MDPV) and α-pyrrolidinopentiophenone (α-PVP) are monoamine transporter inhibitors with high selectivity for the dopamine transporter over the serotonin or norepinephrine transporters (Baumann et al. 2013; Marusich et al. 2014). These compounds are highly effective and potent (e.g. in comparison with methamphetamine or methylone) reinforcers in male rats when assessed with the intravenous self-administration (IVSA) model (Aarde et al. 2013; Schindler et al. 2016; Watterson et al. 2014), and MDPV and α-PVP have been shown to be approximately equipotent when directly compared under a Progressive Ratio schedule of reinforcement in IVSA (Aarde et al. 2015a; Gannon et al. 2017a). A binge-like acquisition pattern of MDPV IVSA has been previously reported for male rats (Aarde et al. 2015b) in a study which also contrasted the effect of concurrent access to an activity wheel. In this pattern, a session was observed early in acquisition for about half of the individual rats wherein drug intake was particularly high on one day, but this was followed by a subsequent pattern of lower, but sustained intake for subsequent sessions. This binge-like pattern was independent of concurrent wheel-access since it occurred in equal proportions of the wheel-access groups. The pattern has not been reported for the acquisition of IVSA of other psychostimulants; however it is unclear if such individual patterns occurred but were overlooked in typical group-mean analysis or if such patterns do not typically occur with other drugs. The present study was therefore designed to determine if the binge-like acquisition would occur during acquisition of α-PVP IVSA, i.e., if the phenomenon would *generalize* to another closely related (see **Figure 1**) drug.

**Figure 1:**
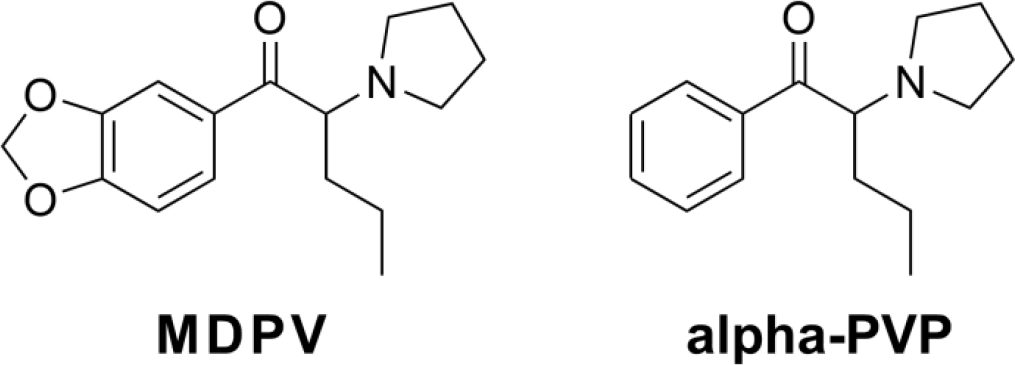
Chemical structures of MDPV and α-PVP.

The prior study of MDPV IVSA included an examination of the effect of concurrent access to an activity wheel because of prior findings that availability of an activity wheel in the home cage reduced IVSA of cocaine, methamphetamine, MDMA or mephedrone on the following session (Aarde et al. 2015c; Smith and Witte 2012), that wheel access provided concurrently with the IVSA session reduced intake of methamphetamine in male rats previously trained to press the lever for food reward (Miller et al. 2012) and reduced cocaine IVSA in female rats with previous wheel experience (Cosgrove et al. 2002). In our MDPV study, concurrent wheel access did not affect the acquisition of MDPV IVSA in behaviorally naïve male rats (Aarde et al. 2015b), however the ongoing IVSA of MDPV was accompanied by a gradual group mean decrease in wheel activity; this was similar to effects observed in male Wistar and Sprague-Dawley rats allowed to IVSA methamphetamine (Miller et al. 2012). Furthermore, the group mean trend for the MDPV study obscured the essentially immediate transition within-animal whereby IVSA of MDPV supplanted wheel activity within a single session for many of the subjects. One secondary goal of this study was therefore to determine if a similar co-option of wheel reinforcement would generalize to rats trained to IVSA α-PVP.

Female rats were used for this study as a further test of the generality of the binge-acquisition phenomenon and to address a general dearth of information on the self-administration of α-PVP, MDPV or indeed most cathinones in female animals (Gannon et al. 2017b; Hadlock et al. 2011; Schindler et al. 2016; Vendruscolo et al. 2017; Watterson et al. 2014). Female rats will self-administer more cocaine (Roth and Carroll 2004b; Smith et al. 2011) and more methamphetamine (Reichel et al. 2012; Roth and Carroll 2004a) than male rats (see (Anker and Carroll 2011; Lynch 2017) for reviews), however the IVSA of entactogen stimulants such as mephedrone, methylone and 3,4-methylenedioxymethamphetamine (MDMA) does not differ substantially between male and female rats (Creehan et al. 2015; Vandewater et al. 2015). Other studies have failed to find any sex-difference in IVSA of cocaine in rats (Miller et al. 2017; Perry et al. 2013) or the oral self-administration of cocaine in mice (DePoy et al. 2016) and in some cases male rats may acquire cocaine IVSA more rapidly and in higher proportion compared with female rats (Swalve et al. 2016a). These findings caution against simple assumptions about sex differences in drug IVSA and recommend investigation of the abuse liability of novel drugs in female as well as male rats to establish similarities and differences that might exist.

This study was also planned to examine any association of the binge-like acquisition phenotype with reinforcer potency or efficacy after acquisition. Some of the popular media reports of disturbing behavior (Alvarez et al. 2012; Anderson 2015; D’Oench 2015; Milian 2015) and medical emergency/fatality (Antonowicz et al. 2011; Benzie et al. 2011; Borek and Holstege 2012; Penders and Gestring 2011; Spiller et al. 2011; Umebachi et al. 2016; Wurita et al. 2014) subsequent to MDPV or αPVP use appear to be associated with binge-like use. Our prior study found that stable post-acquisition MDPV IVSA was higher in the binge-acquisition animals compared with the non-bingers, suggesting either a pre-existing liability or a change caused by the binge-like acquisition. Therefore a dose-substitution experiment was included to determine if α-PVP was more potent or efficacious in the binge-acquisition individuals. A final comparison of the IVSA of α-PVP with MDPV was included to further facilitate inference between the present study and our prior work with male rats.

## Method

### Subjects

Female (N=40) Wistar rats (Charles River, New York) entered the laboratory at 10 weeks of age and were housed in humidity and temperature-controlled (23±1 °C) vivaria on 12:12 hour light:dark cycles. Animals had ad libitum access to food and water in their home cages and remained gonadally intact for this study. All experimental procedures took place in scotophase and were conducted under protocols approved by the Institutional Care and Use Committees of The Scripps Research Institute and in a manner consistent with the Guide for the Care and Use of Laboratory Animals (National Research Council (U.S.). Committee for the Update of the Guide for the Care and Use of Laboratory Animals. et al. 2011).

### Drugs

The α-pyrrolidinopentiophenone HCl (α-PVP) was obtained from Cayman Chemical (Ann Arbor, MI) and 3,4-methylenedioxypyrovalerone HCl (MDPV) was obtained from Fox Chase Chemical Diversity Center (Doylestown, PA). Drugs were dissolved in physiological saline for the i.v. route of administration and all doses are expressed as the salt.

### Intravenous catheter implantation

Rats were anesthetized with an isoflurane/oxygen vapor mixture (isoflurane 5 % induction, 1-3 % maintenance) and prepared with chronic intravenous catheters as described previously (Aarde et al. 2015a; Aarde et al. 2013; Miller et al. 2013; Nguyen et al. 2017). Briefly, the catheters consisted of a 14-cm length polyurethane based tubing (MicroRenathane®, Braintree Scientific, Inc, Braintree MA, USA) fitted to a guide cannula (Plastics one, Roanoke, VA) curved at an angle and encased in dental cement anchored to an ~3-cm circle of durable mesh. Catheter tubing was passed subcutaneously from the animal’s back to the right jugular vein. Catheter tubing was inserted into the vein and secured gently with suture thread. A liquid tissue adhesive was used to close the incisions (3M™ Vetbond™ Tissue Adhesive; 1469S B). A minimum of 4 days was allowed for surgical recovery prior to starting an experiment. For the First 3 days of the recovery period, an antibiotic (cephazolin) and an analgesic (Flunixin) were administered daily. During testing and training, intra-venous catheters were Flushed with ~0 .2–0.3 ml heparinized (32.3 USP/ml) saline before sessions and ~0.2–0.3 ml heparinized saline containing cephazolin (100 mg/ml) after sessions. Catheter patency was assessed once a week, beginning in the third week of training, via administration through the catheter of ~0.2 ml (10 mg/ml) of the ultra-short-acting barbiturate anesthetic, Brevital sodium (1 % methohexital sodium; Eli Lilly, Indianapolis, IN). Animals with patent catheters exhibit prominent signs of anesthesia (pronounced loss of muscle tone) within 3 s after infusion. Animals that failed to display these signs were considered to have faulty catheters and any data that were collected after the previous passing of the test were excluded from analysis.

### Experimental apparatus

Operant conditioning chambers (Med Associates Model ENV-045; Med-PC IV software) enclosed in sound-attenuating cubicles were used for concurrent wheel access and self-administration, as previously described (Aarde et al. 2015b). Chambers were equipped with slanted activity wheels (~100 cm inner circumference, 30 degree tilt of running surface relative to the disk/back), which allowed wheel access while tethered for self-administration. Eight total chambers were available thus animals were tested in daily (M-F) cohorts with N=4 Wheel and N=4 No-Wheel per run.

### Experiments

#### Acquisition and Wheel Switch

Female Wistar rats were prepared with intravenous catheters and trained to self-administer α-PVP (0.05 mg/kg per infusion) using a fixed-ratio 1 (FR1) response contingency in one hour daily (M-F) sessions. A pump pulse calculated to clear non-drug saline through the catheter started the session to ensure the first reinforcer delivery was not diluted, and a single priming infusion was delivered non-contingently if no response was made in the first 30 minutes of the session. The original Wheel group had the wheels unlocked and the No-Wheel group (N=12) had the wheels locked for the first 20 sessions. The wheel-access conditions were then switched for Sessions 21-27. The wheels in two of the experimental chambers were found to be partially malfunctioning to the extent they did not move normally with an animal’s weight and were adjusted on the 6th session. This affected half of the original “Wheel” group (N=12) and created a new Wheel subgroup which effectively had no wheel activity access for the initial 5 sessions. All of the wheels were moveable by rats after this point, however it appeared that this affected self-administration patterns on an individual level. A follow up cohort of 8 Wheel rats was therefore run with N=4 having wheel access from the start of self-administration and N=4 having unlocked wheel access from the 6^th^ session onward to create Wheel and 6-day wheel groups for the group level analysis.

#### Dose Substitution

Following the acquisition and wheel-switch sessions, the rats were subjected to dose substitution (0.0125, 0.025, 0.05, 0.1, 0.3 mg/kg/infusion) sessions in a counter-balanced order (one session per dose). The response contingency was FR1, the sessions length 1 h. Approximately half of the rats first completed a dose-substitution series with the wheel locked and half of the rats with the wheel unlocked (~half of each wheel Group in each wheel condition). The dose-substitution was then repeated with the wheel-access conditions switched such that all rats completed one series with the wheel locked and one series with the wheel unlocked.

#### Estrous Phase

Rats with patent catheters (N=14) were returned to standard training conditions (1 h sessions; 0.05 mg/kg/infusion dose) with the wheels locked for the assessment of any effects of estrous phase. A group of 5 completed the estrous studies with their original catheters intact and a group of 9 had been re-catheterized prior to this study. These studies followed dose-substitution studies with pentedrone and 4-methylpentedrone under FR and PR response contingencies not reported here. Estrous status was assessed via vaginal lavage and cytology using published criteria (Hubscher et al. 2005; Marcondes et al. 2002) after each session (M-F). Self-administration data were assessed over 11-16 consecutive sessions resulting in 1-5 days at each stage for N=14 subjects based on cytology alone. Secondary analysis was conducted on a subset of the data in which a consistent four day progression of the estrous cycle could be identified (N=7; 1-2 sessions per estrous phase).

#### MDPV vs α-PVP

Rats with patent catheters (N=7) were subjected to dose substitution of first MDPV (0.0125, 0.025, 0.05, 0.1, 0.3 mg/kg/infusion) and then α-PVP (0.0125, 0.025, 0.05, 0.1, 0.3 mg/kg/infusion) in a randomized order within drug identity (one session per dose; FR1; session length 1 h; Wheels locked).

### Data Analysis

The number of infusions obtained, wheel activity (quarter rotations) and drug-associated lever discrimination (active lever / all lever responses) in the acquisition and wheel switch sessions were analyzed by Analysis of Variance (ANOVA) with Sessions as a within-subjects (i.e., repeated measures) factor and wheel access Group as a between-subjects factor. Significant main effects were further analyzed with post hoc comparisons of Wheel vs the other groups using the Dunnett procedure. Statistical analysis of wheel activity was limited to the first seven sessions per group, since that was the number of sessions over which the No-Wheel group had access after the wheel-switch. Survival analysis (Mantel-Cox log-rank) was used to compare the rate at which the Groups met an acquisition criterion of 3 sequential sessions with at least 6 infusions obtained; see (Aarde et al. 2015b). A binge was operationally defined for individual rats as a session containing 6 or more infusion in any of the 5 minute intervals. This was derived from a prior work in which non-binge animals obtained less than 5 infusions per 5 minutes during acquisition (Aarde et al. 2015b). The first day on which such a pattern was observed, i.e., binge *acquisition* was the main focus of this analysis, thus it had to occur within the 3 sequential sessions and the rat had to exhibit sustained lower intake after the binge-acquisition session. As in the prior study, any rat that responded for 6 or more infusions in 5 min interval during the apparent acquisition session too late in the session to acquire more total session infusions than in subsequent sessions was assigned the binge phenotype (this affected only #19 in this study). The dose-substitution data were analyzed by ANOVA with Dose and Wheel Access as within-subjects factors. Secondary analysis used Dose as a within-subjects factor and either original training Group or Binge phenotype as a between-subjects factor. Estrous data were analyzed with estrous phase as a within-subjects factor. Significant main effects from these ANOVAs were further analyzed with post hoc multiple comparisons analysis using the Tukey procedure for multi-level factors and the Sidak procedure for two-level factors. Review of the relevant literature (Anker and Carroll 2011; Lynch 2017) supports the hypothesis that drug intake during estrus would be higher than during proestrus thus pre-planned comparisons of estrus versus the other phases was used as the strongest test this hypothesis. The criterion for significant results was at P < 0.05 and all analyses were conducted using Prism 7 for Windows (v. 7.03; GraphPad Software, Inc, San Diego CA).

## Results

### Acquisition

A total of N=25 (N=7 No-Wheel, N=9 Wheel, N=9 6-day wheel) rats retained patent catheters throughout the acquisition and wheel switch parts of the study. Examination of individual patterns of drug self-administration during the acquisition and wheel switch intervals under fixed per-infusion dose conditions confirmed the binge acquisition pattern in 6 / 9 Wheel rats, 5 / 9 6-day Wheel rats (**Figure 2**) and 4 / 7 No-Wheel rats (**Figure 3**). Overall, the three groups of rats self-administered a similar mean numbers of infusions of α-PVP (0.05 mg/kg/inf) during both acquisition and the wheel switch (**Figure 4A**). The rats in the Wheel group with functional wheel access from the start of acquisition were initially delayed in starting IVSA but this was not significantly different and the group ended up with a similar sustained mean intake compared with the other groups. The ANOVA confirmed a significant effect of Sessions of acquisition [F (26, 572) = 7.35; P<0.0001] and of the interaction of factors [F (52, 572) = 1.73; P<0.005], but not of Wheel access Group on infusions obtained. The post-hoc test confirmed that Wheel and 6-day Groups differed only on the final session. The analysis of drug-associated lever discrimination (**Figure 4B**) confirmed only a significant effect of Session [F (26, 572) = 8.75; P<0.0001]. The Mantel-Cox log-rank test failed to confirm (χ^2^ = 3.62, df = 2, P = 0.164) any significant difference between the Groups in terms of the proportion of subjects that reached acquisition criteria during the acquisition interval (**Figure 4D**). Comparison of the first seven sessions of wheel activity across groups (**Figure 4C**) confirmed a significant effect of Group [F (2, 23) = 7.31; P<0.005] and of Session [F (6, 138) = 3.28; P<0.005], but not of the interaction of factors. The post-hoc test confirmed that activity was lower in the No-Wheel and 6-day Wheel groups compared with the Wheel group. Since activity in the No-Wheel group was low after they gained access, and they were being trained in chambers distinct from the original Wheel Group, an additional experiment was conducted wherein a different group (N=8) of female Wistar rats were placed in the set of 8 experimental chambers with unlocked wheels for four one-hour sessions. The rats were in different boxes each session, thus data for each wheel were derived from 4 different rats. This study confirmed that the wheels in the chambers of the No-Wheel group were not differentially functional, i.e., the test group produced similar mean activity levels across all chambers. In addition, the No-Wheel chamber producing the lowest mean activity in this validation test was the one used for the No-Wheel rat that exhibited the most activity of that group during the wheel switch sessions.

**Figure 2:**
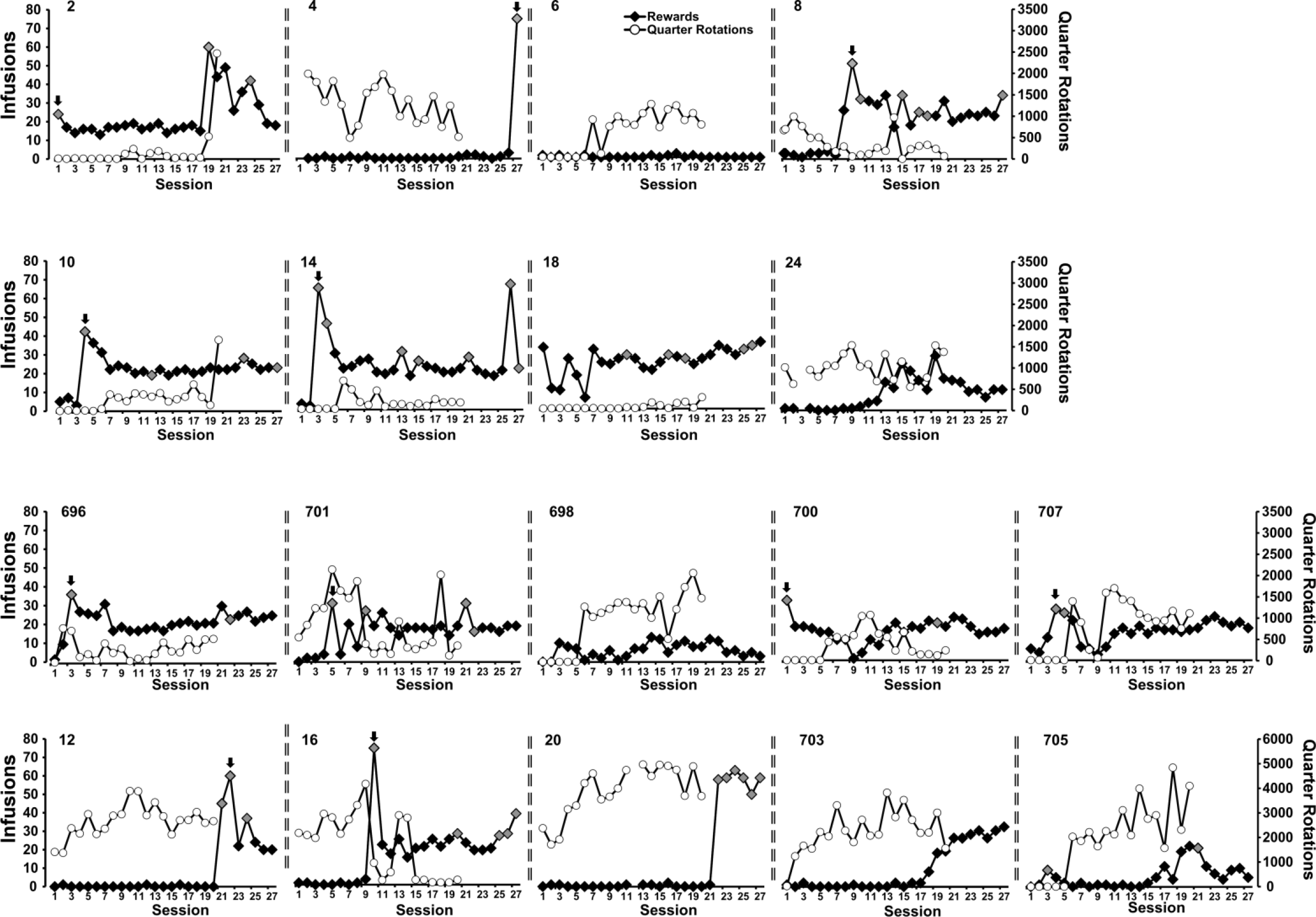
Individual infusions and wheel activity across the acquisition and wheel switch intervals for the Wheel (4, 8, 12, 16, 20, 24, 696, 701, 703) and 6-day wheel (2, 6, 10, 12, 14, 18, 698, 700, 705, 707) groups. Grey symbols highlight the sessions in which 6 or more infusions were obtained in a five-minute interval and the arrow points to the binge-like acquisition sessions.

**Figure 3:**
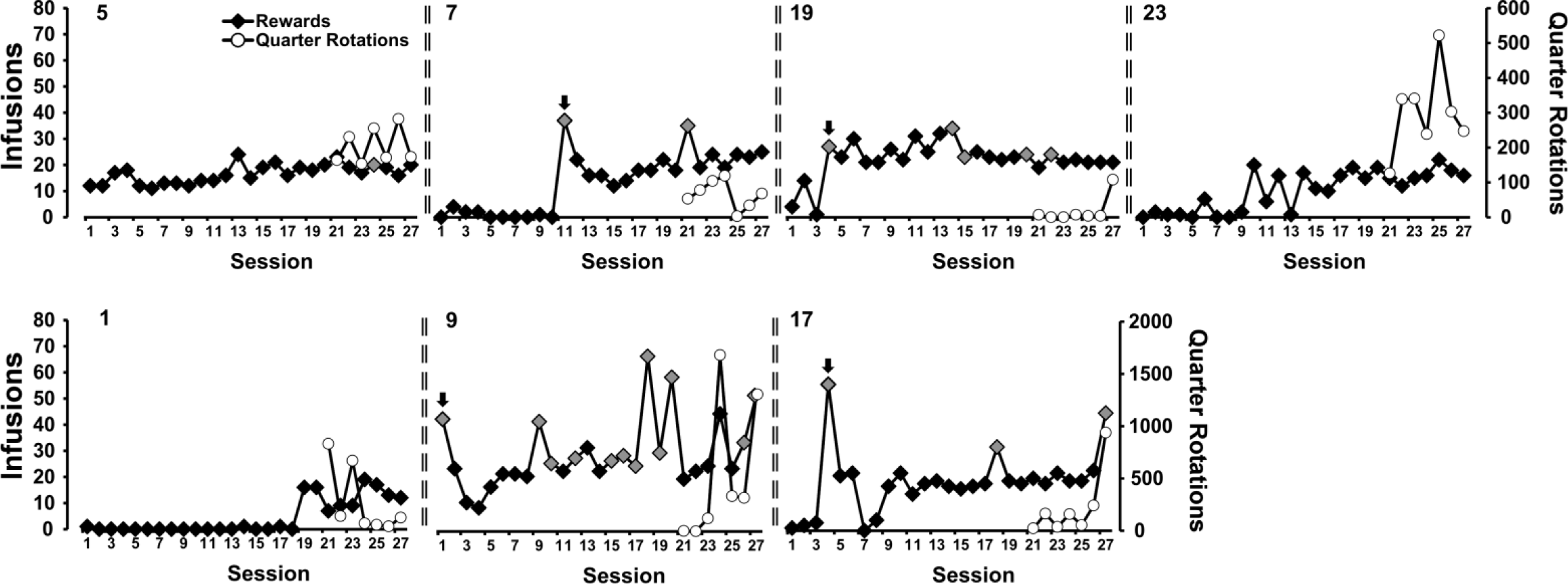
Individual infusions and wheel activity across the acquisition and wheel switch intervals for the No-Wheel acquisition group. Grey symbols highlight the sessions in which 6 or more infusions were obtained in a five-minute interval and the arrow points to the binge-like acquisition sessions.

**Figure 4:**
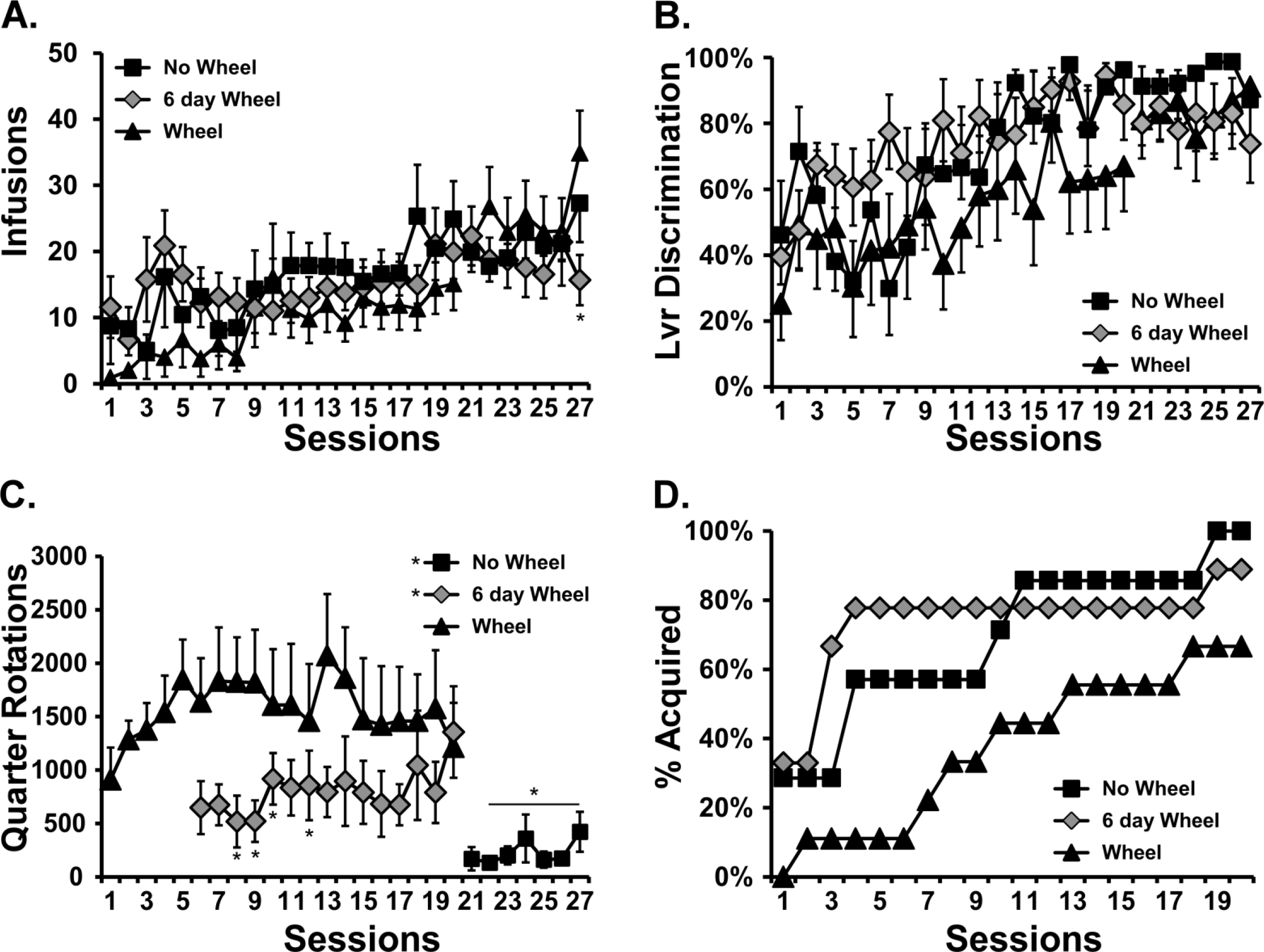
Mean (±SEM) A) infusions, B) percent drug-associated lever responses, C) wheel activity and D) Percent of each group meeting acquisition criteria for Wheel (N=9), 6-day wheel (N=9) and No-Wheel (N=7) acquisition groups. A significant difference from the Wheel group is indicated by *.

The Wheel switch produced no significant mean change in infusions obtained by the Wheel or 6-day Wheel groups upon fixing the wheels in place or by the No-Wheel group upon unlocking their wheels. Three of four Wheel/6-day Wheel individuals with essentially no IVSA intake during acquisition started to self-administer drug after the wheels were locked (see **Figure 2**) but this produced no significant group mean increase.

### Dose-Substitution

#### Self-Administration

A total of N=20 (N=7 No-Wheel, N=7 6-day Wheel; N=6 Wheel) rats retained patent catheters throughout the α-PVP dose-substitution studies. Of these, N=11 exhibited a Binge-like phenotype during acquisition/wheel swap and N=10 did not. Wheel access had no effect on the dose-effect function for α-PVP IVSA in the entire group (**Figure 5A**). The ANOVA confirmed a significant effect of Dose [F (4, 76) = 106.1; P<0.0001] but not of Wheel Access condition or the interaction of factors. The post-hoc test confirmed that each dose produced significantly different numbers of infusions compared with each of the other doses, collapsed across wheel-access condition. There was no effect of acquisition Group during the dose-substitution (**Figure 5B**). The ANOVA confirmed only a significant effect of dose on infusions obtained [F (4, 136) = 159.1; P<0.0001]. Similarly, there was no effect of Binge (N=11) / No Binge (N=10) phenotype on the dose-effect relationship in either Locked or Unlocked wheel conditions (**Figure 5B**). The ANOVA confirmed only a significant effect of Dose [F (4, 132) = 151.8; P<0.0001] on infusions obtained and again, the post-hoc test of the marginal means confirmed that all dose conditions differed from each other.

**Figure 5:**
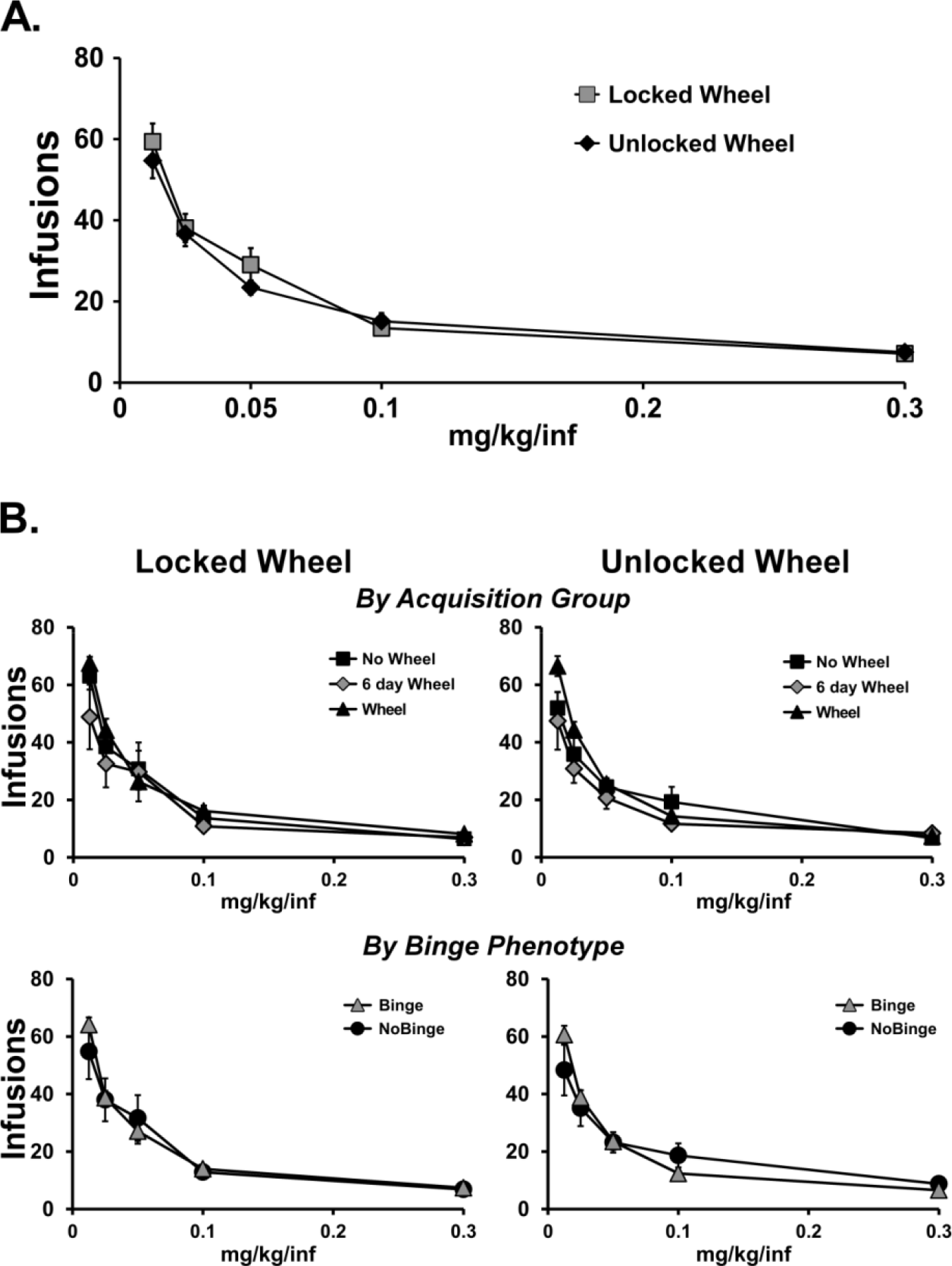
Mean (±SEM) infusions obtained during a dose-substitution. A) All rats with patent catheters (N=20) completed the dose series with the wheel unlocked (Wheel) or locked (No-Wheel). All doses differ significantly from all other doses. B) The results are divided by Acquisition groups of Wheel (N=6), 6 day Wheel (N=7) and No Wheel (N=7) as well as separately by Binge (N=9) and No Binge (N=11) acquisition phenotype subgroups. Data from both subsets are presented for the locked and unlocked wheel dose series.

#### Wheel Activity

Analysis of the Wheel activity during the dose-substitution with the unlocked wheel did not confirm any significant effects of dose, acquisition group or binge phenotype on the number of quarter rotations per session (**Figure 6**).

**Figure 6:**
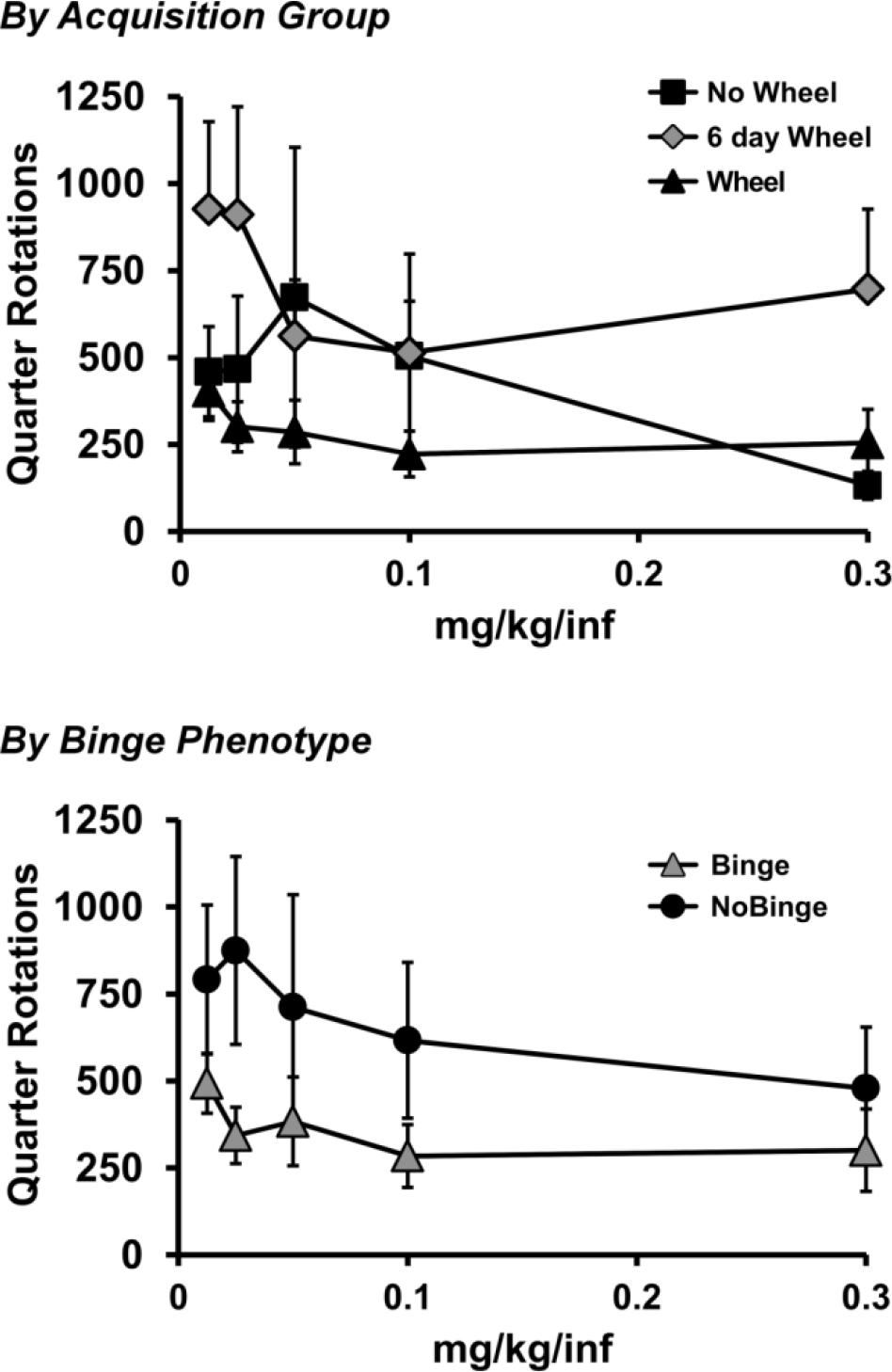
A) Mean wheel activity during the dose-substitution procedure stratified by Acquisition groups of Wheel (N=6), 6 day Wheel (N=7) and No Wheel (N=7) as well as separately by Binge (N=9) and No Binge (N=11) acquisition phenotype.

### Estrous Phase

A total of N=5 rats retained their original catheter patency through the estrous test. An additional N=9 were re-catheterized prior to this phase. There were no differences in the self-administration of α-PVP (0.05 mg/kg/infusion) across the estrous phases (**Figure 7**) and statistical analysis of the full dataset (N=14) confirmed no effects of estrous phase on infusions obtained [F (1.782, 23.16) = 1.12; P=0.34]. Analysis of the subset (N=7) with cycle verified by both post-session cytology and observation of a standard four-day cycle likewise found no effect of estrous stage [F (1.634, 9.802) = 0.41; P=0.63].

**Figure 7:**
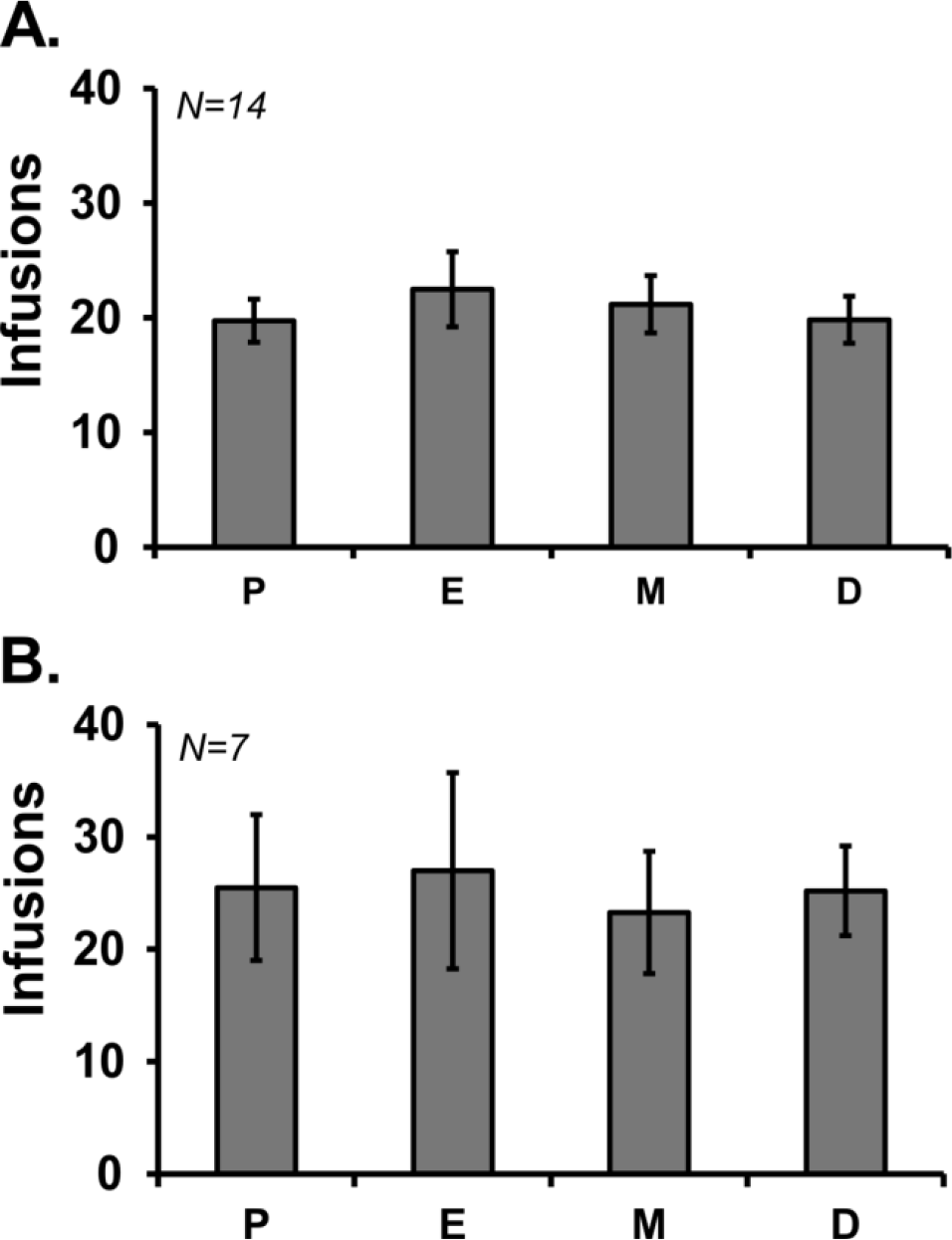
Mean (±SEM) infusions obtained across estrous phases for A) the full sample of 14 rats assessed for estrous effects, or B) the subset of 7 rats with standard 4 day cycles verified. P, proestrus; E, estrus; M, metestrus; D, diestrus.

### MDPV vs α-PVP

A total of N=7 rats retained patent catheters through a final dose substitution contrasting MDPV with α-PVP (**Figure 8**). The ANOVA confirmed a significant effect of Dose [F (4, 24) = 23.18; P<0.0001] and an interaction of drug identity with Dose [F (4, 24) = 3.05; P<0.05]. The post-hoc test confirmed that significantly fewer infusions of MDPV were obtained relative to α-PVP when 0.0125 or 0.25 mg/kg were available per infusion.

**Figure 8:**
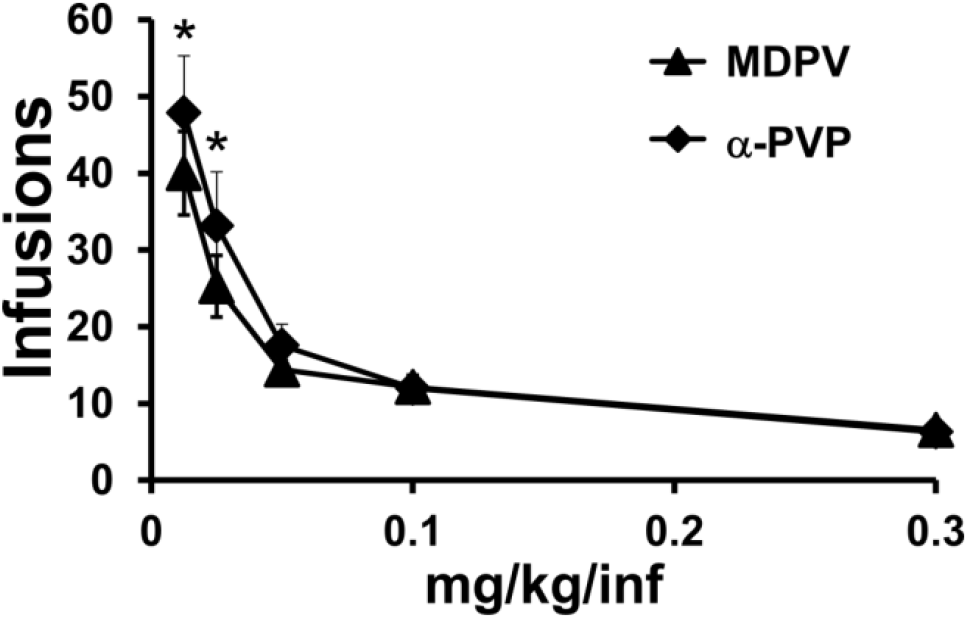
Mean (N=7; ±SEM) infusions of MDPV and α-PVP obtained during the final dose-substitution. A significant difference between compounds is indicated with *.

## Discussion

This study confirms that α-PVP is a highly effective reinforcer in female rats, just as has been previously shown in male rats (Aarde et al. 2015a; Gannon et al. 2017a). The potency and efficacy of α-PVP and the related drug MDPV (Figure 8) were similar in intravenous self-administration (IVSA) in female rats, just as in male rats (Aarde et al., 2015a). Mean cumulative drug intake after 13 sessions of acquisition was similar to that found in a single prior α-PVP IVSA study in which female rats were trained for 13 acquisition sessions (Javadi-Paydar et al. 2017). The amount of α-PVP that was self-administered by the female rats by the end of acquisition in this study was also very similar to the amount obtained by males in a prior study of α-PVP IVSA using similar methods (Aarde et al. 2015a), thus supporting an inference of minimal sex-differences in the IVSA of α-PVP. As with a prior study of MDPV IVSA in male rats (Aarde et al. 2015b), the acquisition of α-PVP IVSA in this study was not altered binge-like session by concurrent access to an activity wheel on a group mean basis. Correspondingly, the introduction of wheel access only after the initial 20 session acquisition period did not decrease IVSA, similar to a prior result for methamphetamine IVSA (Miller et al. 2012). In addition, about half of the rats exhibited a during acquisition after which intake subsided to a stable level that was higher than that observed prior to the binge day. As such, this study shows that the binge-like pattern of acquisition that was previously reported for MDPV IVSA in male rats (Aarde et al. 2015b) generalizes to female rats self-administering α-PVP. Generalization across two variables is a stronger indication of the robustness of the phenomenon compared with generalization across a single variable. The binge session was observed in about half of the Wheel and No-Wheel groups in the prior MDPV study (Aarde et al. 2015b) and this was matched by the relative proportions of binge-like acquisition female rats in the three wheel-access training groups in this study, further supporting the conclusion that this binging pattern is independent of wheel access during training.

The dose-substitution data indicated first that there were no lasting consequences of the binge-like acquisition phenotype, compared with the animals that did not express this phenotype. This shows that the binge is not likely a marker for a priori individual-preference differences that would have significant lasting effects on the potency or efficacy of α-PVP in IVSA. Similarly, this lack of any difference in the dose-substitution study suggests that the day or days of peak brain exposure to α-PVP during the binge did not induce any lasting neuronal plasticity related to rewarding value. Although data for α-PVP are not available, repeated non-contingent dosing with MDPV does not induce significant changes in brain monoamine function (Angoa-Perez et al. 2017; Anneken et al. 2015; Miner et al. 2017) as is the case for amphetamine derivative transporter substrates / monoamine releasers such as methamphetamine and 3,4-methylenedioxymethamphetamine (Green et al. 2003; McFadden et al. 2011; Ricaurte and McCann 1992; Sarkar and Schmued 2010; Taffe et al. 2002). Interestingly a recent study found that five repeated 96 hour MDPV self-administration sessions separated by 72 hours produced only subtle cognitive changes in rats (Sewalia et al. 2017). The interrogation of lasting consequences of the binge acquisition was a major goal of this study as an extension beyond the prior study of male rats self-administering MDPV, however since potential alteration of the dose-response function was not assessed in that prior work it is not possible to compare outcomes on this dimension.

Originally undetected issues with the movement of wheels in two of the chambers used for Wheel access animals and inclusion of additional animals permitted an assessment of a group mean reduction in wheel activity that was observed, which was similar to that assessment in prior studies of MDPV and methamphetamine IVSA in male rats (Aarde et al. 2015b; Miller et al. 2012). The wheel activity of the No-Wheel group in this study after the wheel switch, and in the 6 day wheel group, was lower than the initial activity in the group with functional wheels from the beginning of training. The finding that wheel activity declined in concert with the amount of IVSA experience prior to unfettered wheel access is similar to what was observed in the finding of Miller et al. (2012). The fact that the female rats exhibited about twice as much spontaneous wheel activity than did the prior male group of Wistar rats (Aarde et al. 2015b) emphasizes the phenomenon involves the competition of alternate reinforcers, rather than physical activity as measured by distance-traveled, in this model. This is a further expansion of the finding that male rats strains with a six-fold difference in wheel activity (Wistar > Sprague-Dawley) exhibited a similar degree of suppression of methamphetamine IVSA (Miller et al. 2012) and further underlines the phenomenon by which drug IVSA co-opts and devalues more naturalistic sources of reinforcement.

This study failed to find any effect of wheel access on established IVSA in the switched condition for the original No-Wheel group or the introduction of wheel access in the 6 day group, as well as in the dose-effect functions generated with and without wheel access during the dose-substitution study. This outcome contrasts with the foundational report of Cosgrove and colleagues in which cocaine IVSA and wheel activity were mutually inhibited (Cosgrove et al. 2002). The present finding confirms that our prior finding in which wheel access had no effect on established MA IVSA (Miller et al. 2012) was not a consequence of study in male rats. This is a critical confirmation given that the effect of wheel on cocaine IVSA in Cosgrove et al. (2002) was present in female rats to a much greater extent than in male rats. This study also shows the result is not likely attributable to pharmacological class, since α-PVP and cocaine are both restricted transporter inhibitors whereas methamphetamine is a transporter substrate and releaser.

There is one caveat to the overall conclusion that wheel access had minimal mean effect on the IVSA of α-PVP. Despite the fact that discontinuing wheel access had no significant mean effect on IVSA in the Wheel group (as in our prior study of methamphetamine IVSA), it was the case that three of four individuals who sustained high levels of wheel activity and minimal IVSA behavior during the 20 session acquisition interval started to respond for drug when the wheel was subsequently fixed in place. Thus it may be the case that a subset of rats would fail to ever acquire IVSA if wheel access was always available as an alternative.

This is one of the first available studies of the IVSA of the restricted transporter inhibitor cathinone derivatives in female rats and post-acquisition behavior was both highly stable under a fixed per-infusion dose and parametrically sensitive to dose-substitution. Thus the model offered the opportunity to determine potential effects of estrous phase since some prior reports indicate that cocaine and methamphetamine IVSA change significantly in female rats across the cycle, as reviewed previously (Anker and Carroll 2011; Lynch 2017). No differences in IVSA were found across estrus phases in this study whether phase was determined solely by cytology or when the analysis was restricted to only a subset of animals for which a 4-day cycle could be clearly established. Additional study will be required to determine if there is something unique about α-PVP relative to either cocaine or methamphetamine with respect to differences in self-administration caused by hormonal fluctuation in female rats. Alternately, it is possible that our choice to run animals in the dark cycle, contrasted with prior models which run animals in the light (inactive) part of the day (Lynch et al. 2000; Lynch and Carroll 1999; Ramoa et al. 2013; Smethells et al. 2016; Swalve et al. 2016b), may explain the observed differences.

In conclusion, this study further confirms the likely high abuse liability of the second-generation cathinone α-PVP. The establishment of IVSA by female rats was similar to that of α-PVP or MDPV IVSA in male rats reported in prior studies, including a similar cumulative intake, a reduction in the value of wheel activity as an alternate reinforcer and a binge-like initial acquisition session in about half of the rats. Nevertheless, this study went on to show that the binge-like acquisition phenotype did not have lasting consequences for self-administration behavior, thus it may be the case that the pattern of acquisition is not predictive of individual differences in sustained drug seeking patterns of behavior.

## Acknowledgements

This work was funded by support from the United States Public Health Service National Institutes of Health (R01DA042211) which had no direct input on the design, conduct, analysis or publication of the findings. Conception and securing funding for the project: MAT, TJD; Study design: MAT, MJ-P, JDN; Data collection and initial analysis: MJ-P, ELH, YG, SAV, KMC, JDN; Statistical analysis and figure creation: MAT; Manuscript drafting: MAT, MJ-P. All authors have read and approved the manuscript. The authors declare no competing financial interests. This is manuscript #29574 from The Scripps Research Institute.

